# Disc and Actin-Associated Protein 1 Influence Attachment in the Intestinal Parasite *Giardia lamblia*

**DOI:** 10.1101/2021.08.06.455446

**Authors:** Melissa C. Steele-Ogus, Ava M. Obenaus, Nathan J. Sniadecki, Alexander R. Paredez

## Abstract

The deep-branching eukaryote *Giardia lamblia* is an extracellular parasite that attaches to the host intestine via a microtubule-based structure called the ventral disc. Control of attachment is mediated in part by the movement of two regions of the ventral disc that either permit or exclude the passage of fluid under the disc. Several known disc-associated proteins (DAPs) contribute to disc structure and function, but no force-generating protein has been identified among them. We recently identified several *Giardia* actin (*Gl*Actin) interacting proteins at the ventral disc, which could potentially employ actin polymerization for force generation and disc conformational changes. One of these proteins, Disc and Actin Associated Protein 1 (DAAP1), is highly enriched at the two regions of the disc previously shown to be important for fluid flow during attachment. In this study, we investigate the role of both *Gl*Actin and DAAP1 in ventral disc morphology and function. We confirmed interaction between *Gl*Actin and DAAP1 through coimmunoprecipitation, and used immunofluorescence to localize both proteins throughout the cell cycle and during trophozoite attachment. Similar to other DAPs, the association of DAAP1 with the disc is stable, except during cell division when the disc disassembles. Depletion of *Gl*Actin by translation-blocking antisense morpholinos resulted in both impaired attachment and defects in the ventral disc, indicating that *Gl*Actin contributes to disc-mediated attachment. Depletion of DAAP1 through CRISPR interference resulted in intact discs but impaired attachment, gating, and flow under the disc. As attachment is essential for infection, elucidation of these and other molecular mediators is a promising area for development of new therapeutics against a ubiquitous parasite.

**Author Summary:** *Giardia lamblia* is a single-celled organism and one of the most common gastrointestinal parasites worldwide. In developing countries, recurrent *Giardia* infections are common, due to lack of access to clean water. *Giardia* infections can lead to diarrhea, vomiting, dehydration, disruption of the intestinal microbiome, and chronic infections can lead to irritable bowel syndrome. Because existing drug treatments have side effects and *Giardia’s* resistance to drugs is increasing, new treatment strategies are needed. The parasite’s attachment to the host’s intestine is mediated by a *Giardia*-specific structure that resembles a suction cup and is called the ventral adhesive disc. We previously identified DAAP1, a protein which interacts with *Giardia* actin and localizes to the ventral disc. Here, we explore the relationship between these two proteins and investigate their role in disc-based attachment. Most disc proteins, including DAAP1, are unrelated to any human proteins, making them appealing drug targets to inhibit parasite attachment and infection.

## Introduction

*Giardia lamblia,* a single-celled parasite, is the cause of over 280 million annual cases of the gastrointestinal disease giardiasis worldwide [1,2] This extracellular parasite colonizes the lumen of the intestine, where its ability to maintain infection requires tight attachment to microvilli, in order to withstand the peristaltic flow of the gastrointestinal tract [3]. The primary means of attachment is via a specialized microtubule-based organelle known as the ventral adhesive disc [4–6]. A well-functioning disc is critical for cell survival and disease transmission [7], but the molecular mechanisms controlling conformational dynamics and function remain poorly understood.

Imaging of fluid flow at the disc margin identified two areas postulated to be important for tight attachment: one at the anterior and the other at the posterior of the disc [8–10] (Figure 1A). The former is the marginal region of the overlap zone, the region of the disc where the sheet of microtubules overlaps itself. The other is the ventral groove, a hump in the posterior region of the disc where the ventral flagella exit between the disc and cell body. Interruption of contact in these areas results in disruption of attachment and in fluid moving underneath the disc [9,11,12]. Cryo-electron tomography imaging of the disc reveals different protein densities in these areas, indicating specialized architecture within the disc itself, which may indicate that specific effector proteins control disc conformation and fluid flow [13]. In the current model, both hydrodynamic flow and suction contribute to the mechanism of attachment [9]. Attachment occurs in several distinct stages [8,9]. In early attachment, the anterior ventrolateral flange, lateral crest and ventral groove make contact with the surface while the ventral flagella beat to generate fluid flows. During this stage cells can either skim the surface or remain stationary. In the later stages of attachment, the cell body becomes more intimately associated with the surface so that the lateral shield and bare area make contact with the surface and the cells remain in place. Flagella-beating is required for initiation, but not maintenance, of attachment, indicating that fluid must be moved out from underneath the disc during the earlier stages of attachment [9,14].

**Figure 1:**
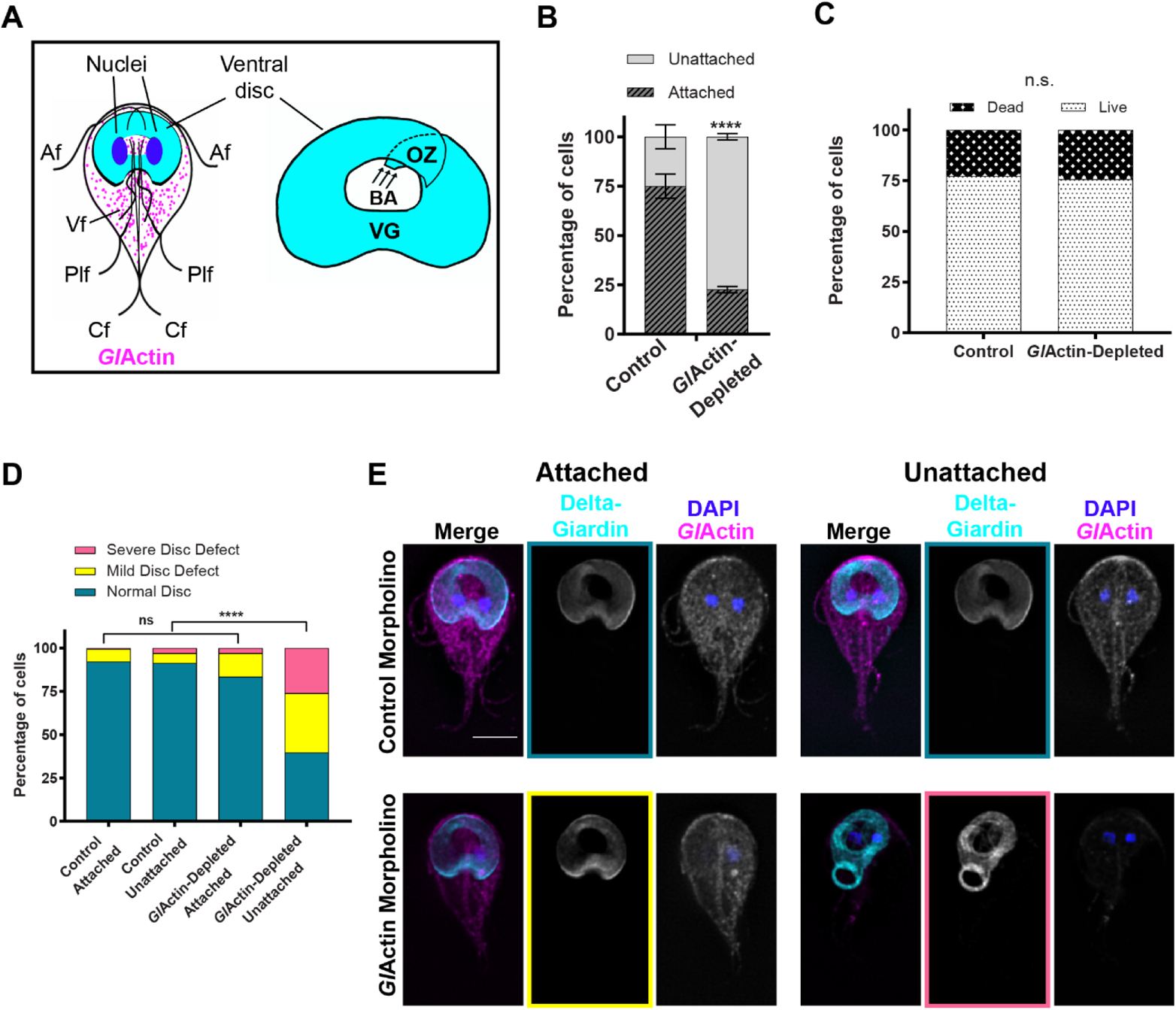
Depletion of *Gl*Actin affects attachment and ventral disc morphology. **A)** Left: Diagram of a *Giardia* trophozoite showing *Gl*Actin (magenta), disc marker delta-giardin (cyan), and DAPI/nuclei (blue). Anterior (Af), posteriolateral (Plf), ventral (Vf), and caudal flagella (Cf) are shown. Right: Enlarged diagram of ventral disc and areas: overlap zone (OZ), ventral groove (VG), and bare area (BA). Arrows indicate the origin of the microtubule sheet and orientation of the microtubule plus ends. **B)** Quantification of non-challenged attachment assays. Attached and unattached cells were counted 24 hours after treatment with either a standard control or anti-*Gl*Actin morpholino. Graph is representative of an average of three replicates, mean ± sem. For control, 74% ± 6 were attached and 25% ± 6 were detached. For *Gl*Actin-depleted cells 23% ± 2 were attached and 77% ± 2 detached. Raw numbers and detailed statistics reported in Supplemental Table 1. Each replicate was analyzed separately with a two-sided chi-square test with Yates’ correction, P<0.0001 for each. **C)** Unattached cells from B were stained with propidium iodide and fluorescein diacetate to distinguish dead from living cells respectively. N=198 control cells and 222 *Gl*Actin-depleted cells. There was no significant difference between the two groups based on a two-sided Fisher’s exact test P= 0.7307. **D)** Both attached and unattached *Gl*Actin-depleted and control cells were fixed and immunostained for delta-giardin to assess ventral disc morphology using fluorescence microscopy. N=126 cells for each condition. There was a significant difference in unattached cells based on a two-sided chi-square test with Yates’ correlation, P<0.0001. No significant difference was detected in attached cells based on the same test, P<0.0903 **E)** Representative images of cells from D, with examples of normal discs (teal), mild defects (yellow), and severe defects (pink). Scale bar denotes 5 µm.

These conformational changes in the disc which control attachment are not generated by microtubule turnover, as microtubule depolymerizing or stabilizing drugs do not affect attachment [15]. Though microtubules are the foundation of the disc, it is also composed of over 100 other proteins, known as disc-associated proteins (DAPs). At least five of these DAPs have been shown to be important for disc biogenesis and function [9,12]. Only three DAPs identified to date contain known microtubule-binding motifs; most are ankyrin-repeat proteins, NEK kinases, or the annexin-like alpha-Giardins [15,16]. At least 25, however, lack identifiable domains and are annotated in the *Giardia* genome as “hypothetical proteins” [17,18]. So far, no DAPs have been identified as motor or force-generating proteins. Thus, the proteins that mediate changes in disc conformation are unknown.

Actin has a conserved role in force generation and in controlling cell and organelle shape in eukaryotic cells[19]. Previously, localization studies using heterologous antibodies demonstrated that *Giardia* actin (*Gl*Actin) and actin regulators associate with the ventral disc [6,20]. This observation led to multiple experiments attempting to test the role of *Gl*Actin in attachment using actin inhibitors. These studies yielded conflicting results because reagents that typically recognize actin or actin-binding proteins in other organisms were presumed to act similarly in *Giardia* [6]. Later, *Gl*Actin was determined to be one of the most divergent actins found among eukaryotes and its genome lacked genes encoding conventional actin-binding proteins that were reportedly localized to the disc [21–23]. Further research revealed that *Gl*Actin has non-conservative amino acid substitutions, predicted to disrupt actin inhibitor binding, and anti-*Gl*Actin antibodies revealed localization throughout the cell [22]. Thus, *Gl*Actin’s potential role in disc-based attachment is controversial.

We recently identified eight *Giardia* actin (*Gl*Actin) interactors that localize to the ventral disc [18], suggesting a means to resolve the role of actin and actin-binding proteins in ventral disc function. One such actin-interacting protein, GL50803_16844, which we denote Disc and Actin Associated Protein 1 (DAAP1), appears highly enriched in the two regions of the disc important for regulating fluid flow during attachment.

In this study, we use morpholinos and CRISPR interference (CRISPRi) to deplete *Gl*Actin and DAAP1 respectively and investigate the role of both proteins in the ventral disc. Our results show that *Gl*Actin plays a role in normal ventral disc morphology and both *Gl*Actin and DAAP1 are required for parasite attachment.

## Results

### *Gl*Actin affects attachment and disc morphology

*In vitro, Giardia* normally grows attached to the inside of a culture tube. We used an antisense translation-blocking morpholino to deplete *Gl*Actin for 24 hours, [22] after which we assessed their attachment. We compared the numbers of cells attached to the tube wall to cells unattached in the media for each condition. With an approximately 50% knockdown (Figure S1), we found attachment to plastic culture tube walls to be significantly decreased, with an average of 23% of cells attached compared to 75% of cells treated with a control morpholino (Figure 1B). Dead *Giardia* cells lose their ability to adhere to culture tubes. To account for the possibility that the majority of the unattached cells were dead, we stained the unattached cells with propidium iodide to mark dead cells, and fluorescein diacetate to mark live cells. While slightly more unattached *Gl*Actin knockdown cells were dead compared to that of the control population, this difference was not statistically significant (Figure 1C), indicating that the defect in attachment in *Gl*Actin-depleted cells was not due to a preponderance of dead cells.

We asked if the role of *Gl*Actin in attachment could be attributed to an effect on the ventral disc. Attached and unattached cells were treated with either control or anti-*Gl*Actin morpholino, fixed after 24 h, and examined for distribution of a ventral disc marker using immunofluorescence microscopy. We used a HALO-tagged delta-giardin which is widely distributed in the disc to monitor whether the disc appeared normal or defective. Defects were then further classified as 1) mild, with misshapen/broken discs or small gaps in the disc or 2) severe, including unwound discs or severely misshapen discs (Figure S2, 1D, 1E). More *Gl*Actin knockdown cells had defective discs compared to those in the control; however, this difference was only significant in unattached cells (Figure 1D, 1E). While both attached and unattached *Gl*Actin-depleted cells had more disc defects than their counterparts treated with the control morpholino, the unattached *Gl*Actin-depleted cells had the most defective discs among all the categories. Yet, there were more unattached cells than cells with disc defects, suggesting that *Gl*Actin has a role in attachment beyond its contribution to disc morphology.

### *Gl*Actin interacts with disc-associated protein DAAP1

We next asked if GL50803_16844, previously identified by mass spectrometry as a potential actin filament interactor at the ventral disc [18], could be involved in attachment. GL50803_16844 is conserved in all *Giardia lamblia* isolates and *Giardia muris*, but is absent from the genome of *Spironucleus*, a close relative of *Giardia* which lacks a ventral disc [24]. Hereafter, we will refer to this protein that lacks identifiable domains and homologues in other organisms as Disc- and Actin-Associated Protein 1 (DAAP1). We tagged DAAP1 with a C-terminal 3xHA tag and expressed it episomally in *Giardia* (Figure S2). Immunoprecipitation of this construct revealed it in complex with *Gl*Actin (Figure 2A). We also endogenously tagged DAAP1 with a C-terminal mNeonGreen tag then used live and fixed cell fluorescent imaging. Notably, this fusion protein localizes to the entirety of the disc, but it is highly enriched in the ventral groove and the overlap zone, two areas implicated in attachment (Figure 1A, 2B).

**Figure 2:**
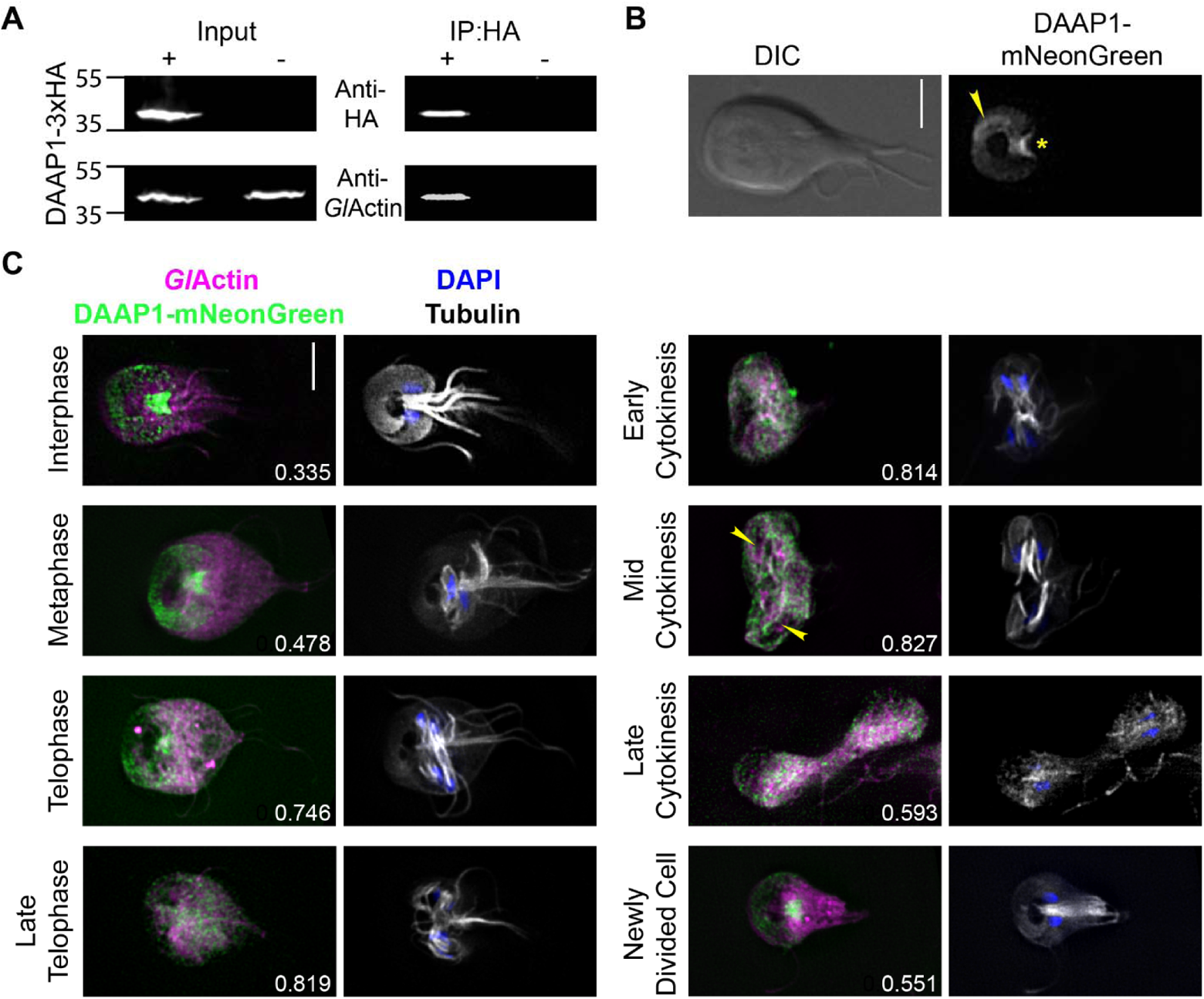
DAAP1 is a *Gl*Actin interactor and a ventral disc-associated protein. **A)** Immunoprecipitation from extracts of cells expressing DAAP1-3XHA and wild type cells, followed by western blots probed with anti-*Gl*Actin and anti-HA antibodies, revealed interaction between the two proteins. **B)** Image of DAAP1-mNeonGreen in a live cell. The fusion protein localizes to the disc and is enriched in the ventral groove (asterisk) and overlap zone (arrow). Scale bar denotes 5 µm. **C)** Immunofluorescence of *Gl*Actin (magenta), tubulin (cyan), DAAP1-mNeonGreen (green), and DAPI/nuclei (blue) during the cell cycle. DAAP1-mNeonGreenremains on the disc into telophase, is incorporated into the nascent discs by mid cytokinesis (see arrows), and localization is fully restored in newly divided cells. Pearson’s correlation coefficients using Costes randomization were calculated for colocalization between DAAP1 and *Gl*Actin in each image, shown in the lower righthand corner. The Costes P-value was 1 for all images. Scale bar denotes 5 µm.

Flexion of the overlap zone can be seen during detachment from the surface, consistent with the notion that conformational changes of specific regions of the disc can regulate attachment (Supplemental Movie 1).

To investigate co-localization between *Gl*Actin and DAAP1, we calculated Pearson’s correlation coefficients for these two proteins using the JACoP plugin for ImageJ [25,26]. While DAAP1-mNeonGreen does not appear to co-localize with *Gl*Actin during interphase, in mitosis DAAP1 leaves the ventral disc and was observed to co-localize with *Gl*Actin (Figure 2C).

We also measured the intensity of DAAP1-mNeonGreen fluorescence in *Gl*Actin-depleted cells, selecting a region in the ventral groove, as that is where DAAP1 expression in the highest (Figure 3). While there was no significant difference between fluorescence levels in attached cells (Figure 3A, 3B), this difference was significant in unattached cells (Figure 3C, 3D). Furthermore, in some *Gl*Actin-depleted cells, we observed DAAP1 loss of enrichment in the entire disc (Figure 3C). These results indicate that *Gl*Actin is responsible for DAAP1 disc localization.

**Figure 3:**
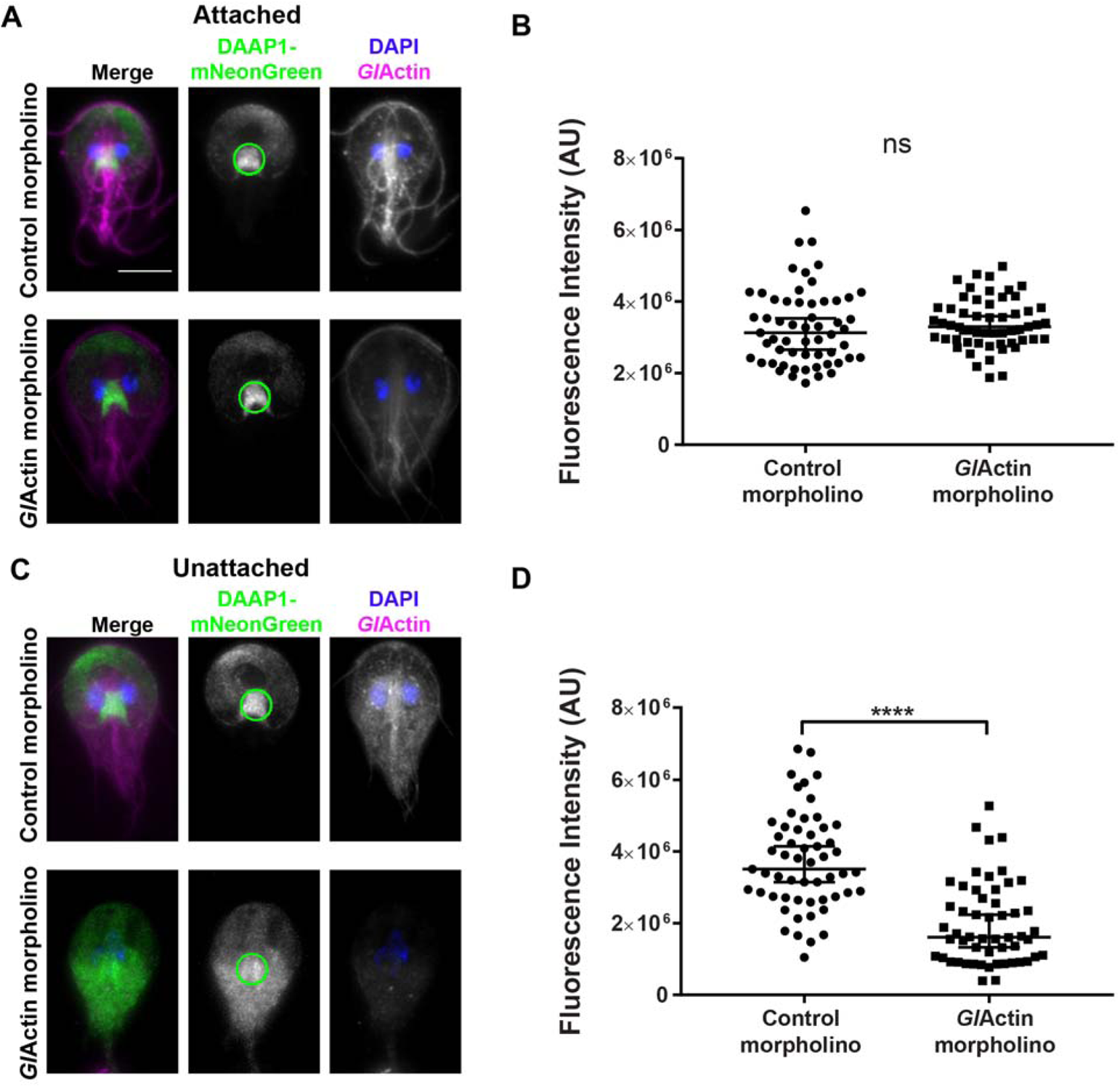
DAAP1 loading onto the ventral disc requires *Gl*Actin. **A)** Representative images of attached cells 24 hours after treatment with either an anti-*Gl*Actin or control morpholino. Fluorescence intensity of DAAP1 at the ventral groove (see green circle) was measured at the brightest section in each stack. *Gl*Actin shown in magenta, mNeonGreen-DAAP1 in green, DAPI in blue. Scale bar denotes 5 µm. **B)** A two-tailed Mann-Whitney test indicated that this difference was not statistically significant, (N_Control_=57, N*_Gl_*_Actin-depletd_=55), P=0.227. Median and 95% confidence interval shown. Median of control=3131445. Median of *Gl*Actin-depleted=3298353. Two and four outliers were detected by Tukey Fence, k=1.5 in the control and *Gl*Actin-depleted condition respectively and were not included in the analysis. **C)** Representative images of unattached cells 24 hours after treatment with either an anti-*Gl*Actin or control morpholino. Fluorescence intensity of DAAP1 at the ventral groove (see green circle) was measured at the brightest optical section in each stack. **D)** A two-tailed Mann-Whitney test indicated that this difference was statistically significant, (N_Control_=55, N*_Gl_*_Actin-depletd_=54), P<0.0001. Median and 95% confidence interval shown. Median of control= 3508064. Median of *Gl*Actin-depleted=1609904. Four outliers were detected by Tukey Fence, k=1.5 in the *Gl*Actin-depleted condition and were not included in the analysis.

The microtubules in the ventral disc are stable and do not undergo dynamic instability; they rely on a set of core DAPs to maintain this stability [15]. During mitosis, the disc is dismantled as the cells must detach to “swim” away from each other during cytokinesis [27,28]. New discs in the daughter cells appear during early cytokinesis and are built from microtubules originating from the mother disc and the median body [27,29]. Notably, DAAP1-mNeonGreen remains on the disc during telophase, late into disassembly, and begins to coalesce again on the daughter discs in mid cytokinesis. In newly-divided cells, DAAP1-mNeonGreen has returned to its disc localization and is once more enriched in the ventral groove (Figure 2C). Fluorescence recovery after photobleaching (FRAP) confirmed the stability of DAAP1-mNeonGreen for disc localization, as fluorescence levels did not recover for 12 minutes after photobleaching (Figure 4) in either of the two regions of the disc where DAAP1 is enriched. Viability was confirmed by cell movement as seen in Supplemental Movie 2. This result is similar to the fluorescence recovery of other DAPs. Taken together, these data indicate that DAAP1 is a *Gl*Actin interactor and stably localized to the ventral disc in interphase.

**Figure 4:**
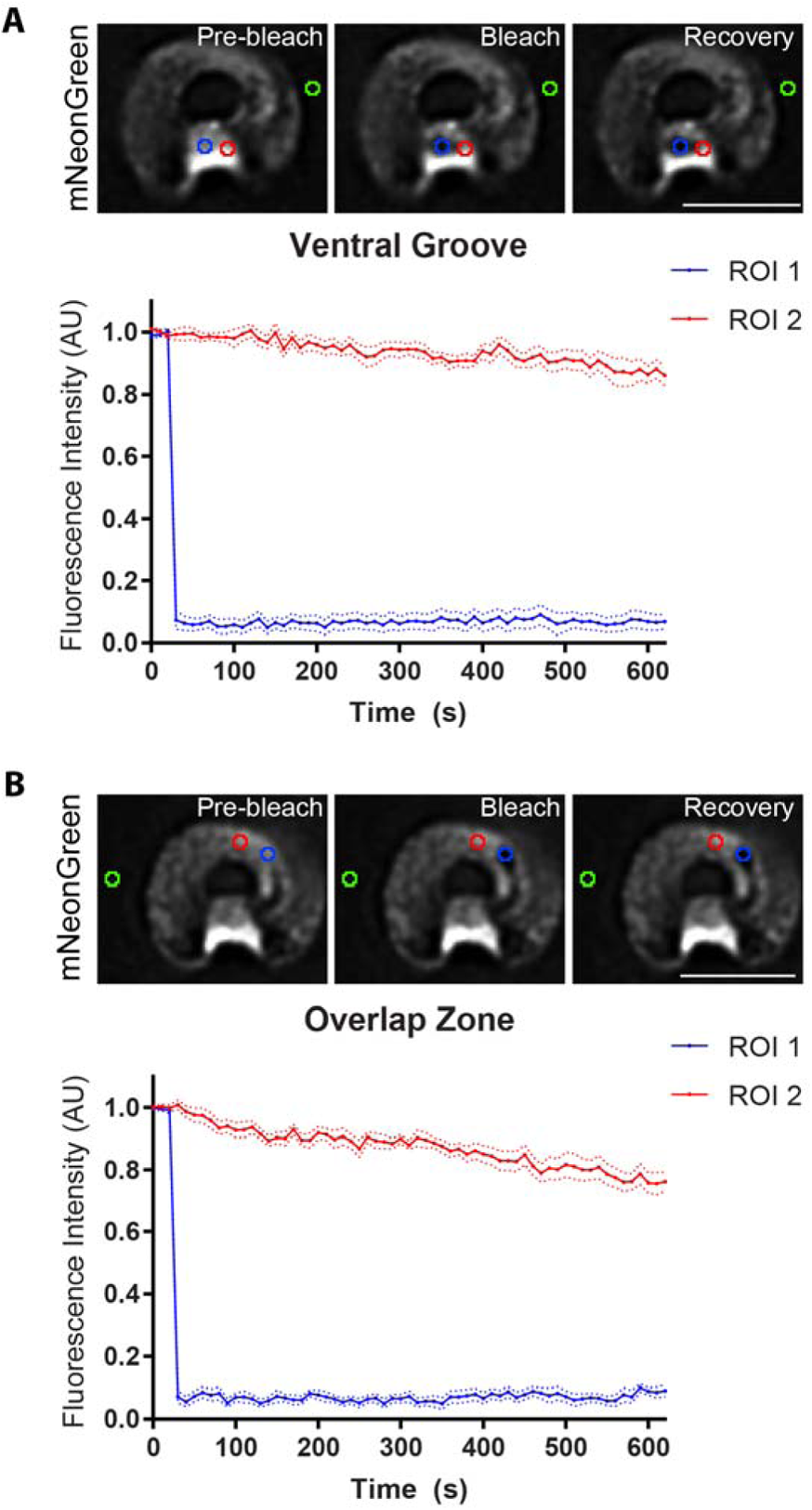
DAAP1 is stably associated with the ventral disc. Fluorescence recovery after-photobleaching (FRAP) in DAAP1-mNeonGreen-cells showed no recovery after 12 minutes for bleached areas in both the ventral groove **(A)** and overlap zone **(B).** Three points of reference were taken, including bleached (ROI1, blue), unbleached (RO2, red), and background (green), and normalized accordingly. Scale bar denotes 5 µm. N=10 for each ROI in both A and B.

### DAAP1-depleted cells have intact ventral discs

Since *Gl*Actin is involved in both attachment and disc morphology, we asked if DAAP1 could have a similar role. We attempted depletion of DAAP1 with translation-blocking morpholinos, however, the sequence was design-resistant and the knockdown level inconsistent. We then turned to CRISRPi to investigate DAAP1’s potential role in attachment and disc morphology. To monitor disc morphology as well as deplete DAAP1, we inserted the HALO-tagged delta-giardin marker into the dead Cas9- (dCas9) based CRISPRi plasmid and introduced this into a DAAP1-mNeonGreen expressing cell line. This allowed us to evaluate knockdown on both a whole population (Figure 5A) and an individual cell basis. We analyzed the fluorescence of endogenously-tagged DAAP1-mNeonGreen in control cells expressing dCas9 with a non-specific guide RNA to those expressing a DAAP1 specific small guide RNA resulting in an average decrease in expression of 40% for the population. We once again separated cells that were attached to culture tubes from unattached cells to probe for disc defects in each subpopulation. We did observe a higher proportion of disc defects in the DAAP1-depleted cells than in the control, but this difference was not significant (Figure 5B and 5C). These results indicate that a 40% depletion of DAAP1 did not impact gross ventral disc morphology, as assessed by light microscopy.

**Figure 5:**
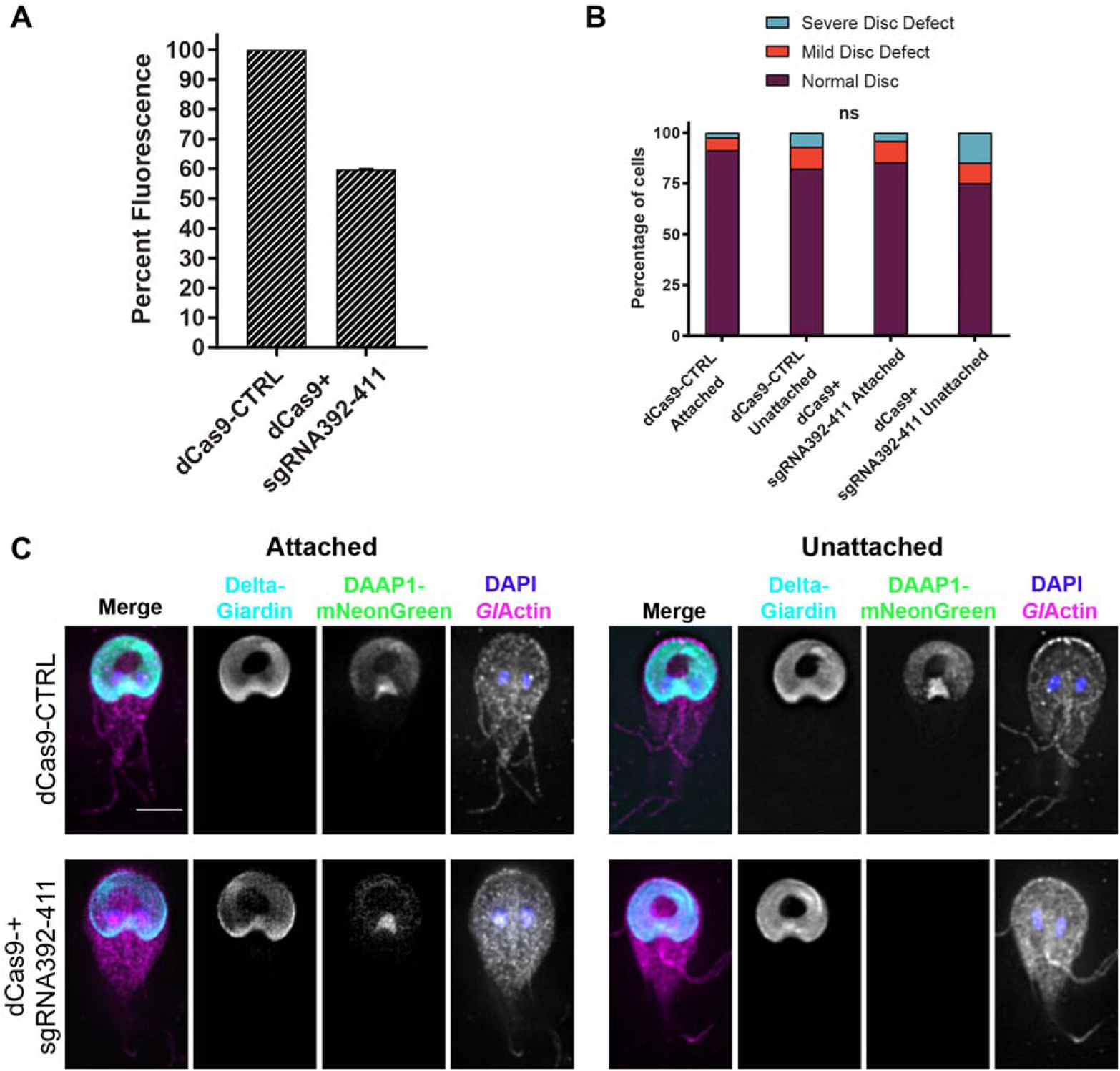
DAAP1-depleted discs do not have defects detectable by light microscopy. **A.)** Fluorescence of DAAP1-mNeonGreen cells transfected with a dead Cas9 vector containing either a non-specific small guide RNA (dCas9-CTRL) or a small guide RNA targeting DAAP1 (dCas9+sgRNA392-411). Normalized to the control, the DAAP1-depleted cells demonstrated an average knockdown of 40% ± 3 (three independent transformations). Attached and unattached control and DAAP1-depleted cells were fixed and stained separately before assessing their disc morphology. Based on a chi-square test, there was no significant difference in disc defects observed between either attached or unattached groups. (Control attached N=129 cells, control unattached N=126, DAAP1-depleted attached N=123, DAAP1-depleted unattached N=128.) Attached cells, P=0.3483.Unattached cells:, P=0.1288. **C.)** Representative images from B: *Gl*Actin (magenta), delta-giardin (cyan), DAAP1 (green), and DAPI/nuclei (blue). Scale bar denotes 5 µm.

### Inability to Maintain Attachment in DAAP1-Depleted Cells

DAAP1 knockdown cells had no visible defects, so DAAP1 may not be needed for disc assembly. To investigate a possible role in attachment, we used the aforementioned attachment assay on DAAP1-depleted cells and observed a small but significant decrease in attachment for two of three replicates relative to control cells (Supplemental Table 2). As CRISPRi is known to have variable penetrance in *Giardia* [30], we analyzed the fluorescence in unattached and attached cells separately. Here we found reduced levels of DAAP1-mNeonGreen in unattached cells compared to attached ones (Figure 6A). This result prompted us to further explore the possible role of DAAP1 in attachment using a shear force attachment assay. In their normal intestinal habitat, *Giardia* must maintain attachment even when subjected to the flow of chyme. With this in mind, we challenged DAAP1-depleted cells with a fluid shear force. DAAP1-depleted and control cells were allowed to attach inside a flow chamber and then subjected to a flow of growth medium. Cell position and velocity were tracked using Trackmate (Figure 6B, [31]). DAAP1-depleted cells slid significantly faster under flow than their control counterparts (Figure 6C, 6D). Furthermore, more control cells remained in the field of view than DAAP1-depleted cells over time (Figure 6E), indicating a better ability to maintain attachment.

**Figure 6:**
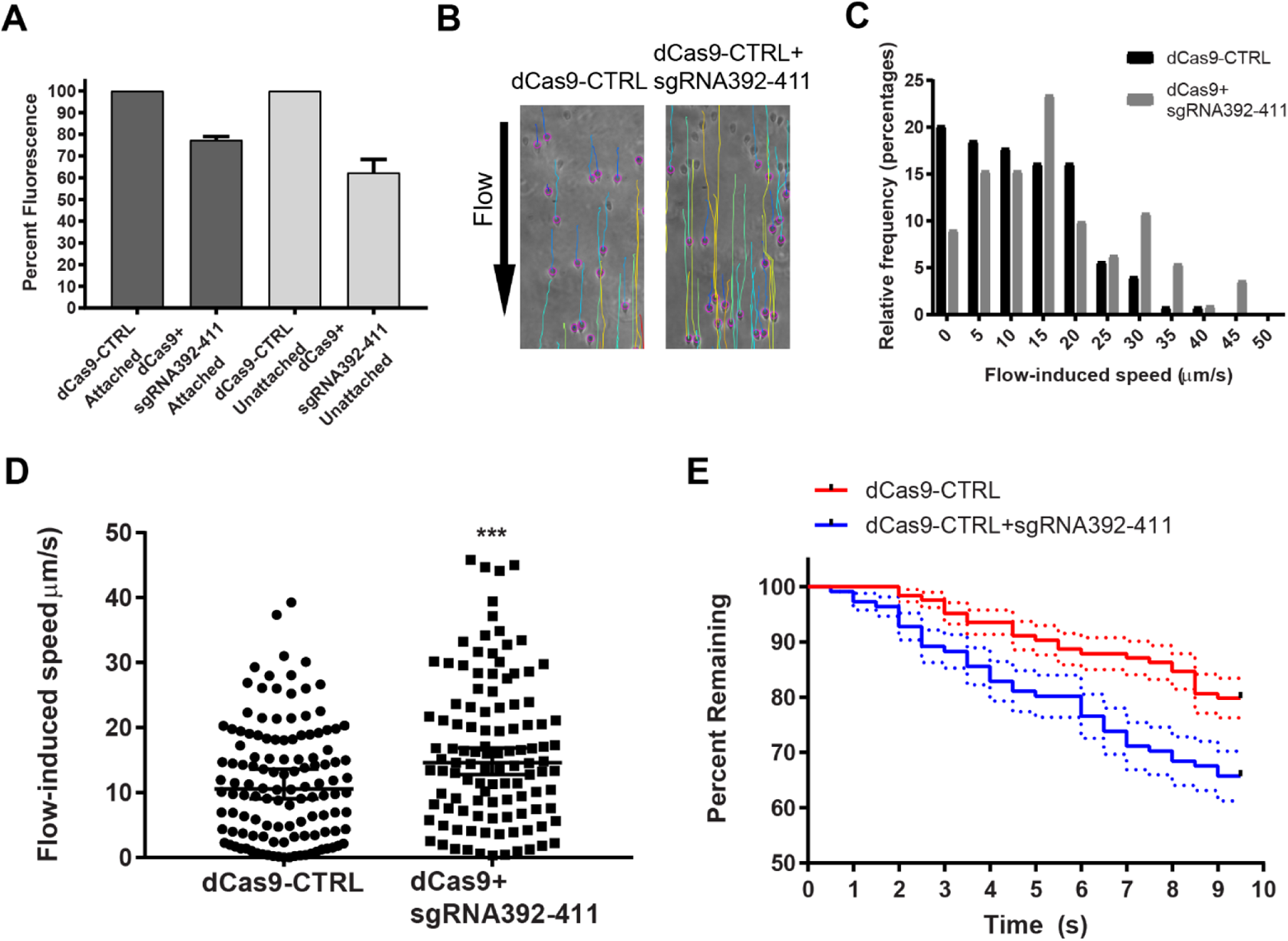
DAAP1 depletion results in reduced attachment. **A)** DAAP1 fluorescence levels from control (dCas9-CTRL) and DAAP1-depleted (dCas9+sgRNA392-411). Unattached cells had a greater decrease in DAAP1 expression (38% ± 6) than attached cells (23% ± 2). Average of two independent transformations with three technical replicates for each is shown. **B)** In this flow chamber assay, attached cells were challenged with a flow rate of 100 µL/min. Cells were tracked for 10 seconds using TrackMate software in ImageJ, colored lines indicating resulting paths. **C)** Distribution of mean flow-induced speed. **D)** Mean flow-induced speed of each cell for 10 seconds after challenge, with median and 95% confidence intervals. A two-tailed Mann-Whitney test indicated that this difference was statistically significant, (N_dCas9-CTRL_=124, N_dCas9+sgRNA392-411_=111), P=0.001. Median of control=10.56, upper 95% CI= 13.33, lower 95% CI=10.15. Median of DAAP1-depleted is 14.62, upper 95% CI=18.62, lower 95% CI=14.41. **E)** Kaplan-Meier curve of cells displaced from the field of view over time, P= 0.0077 N=124 for control, N=111 for DAAP1-depleted cells. Median and standard error shown.

### Impaired Seal Formation in DAAP1-Depleted Cells

We then hypothesized that the impaired attachment in DAAP1-depleted cells may be due to impairments of the ventral groove or overlap zone “gates” that control fluid flow and buildup of negative pressure differentials under the disc. We used fluorescent microspheres to investigate this fluid flow in live, attached cells imaged in normal growth media. We found that spheres entered under the disc more frequently in DAAP1-depleted than dCas9 control cells, consistent with a defect in gating flow under the disc (Figure 7A). Furthermore, we found more beads per cell in the DAAP1-depleted cells (Supplemental Table 4).

**Figure 7:**
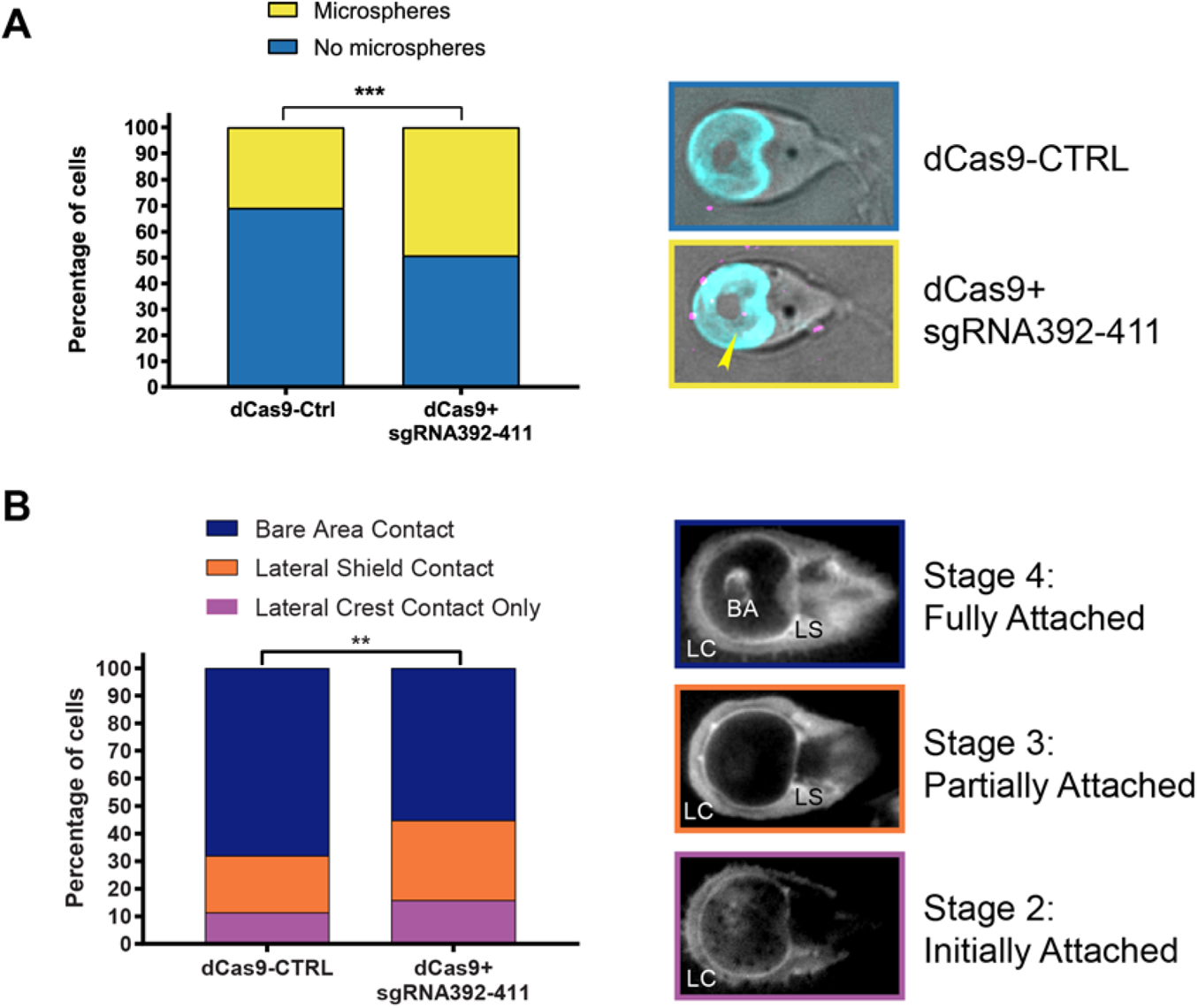
DAAP1 has a role in ventral disc seal formation. **A.)** Fluorescent microspheres were added to live attached control (dCas) and DAAP1-depleted (dCas9+sgRNA392-411) and imaged. Left: significantly more cells with microspheres under the disc were observed in DAAP1-depleted cells based on a chi-square test exact test, P<0.001. N=175 for control, 180 for DAAP1-depleted cells. Right, top: control cell. Right, bottom: DAAP1-depleted cell with microsphere under disc (yellow arrow). Merged images shown with DIC, microspheres (magenta), delta-giardin (cyan). Two-sided Fisher’s exact test, P=0.0156 was used for analysis. **B.)** TIRF microscopy assay to monitor the quality of attachment. Control and DAAP1-depleted cells were stained with the plasma membrane stain CellMask Orange and imaged with TIRF microscopy. Left: significantly more DAAP1-depleted cells were partially attached than control cells based on a chi-square test, N=317 for each condition, P<0.005. Right: images of cells in initial attachment (stage 2) with lateral crest (LC) contact, partial attachment (stage 3), with lateral crest and lateral shield (LS) contact, or fully attached (stage 4), with lateral crest, lateral shield, and bare area (BA) contact.

Since the ability of the gates to close affects disc seal formation [8,14], we used plasma membrane stain, CellMask Orange and Total Internal Fluorescence (TIRF) microscopy to evaluate seal formation for DAAP1-depleted and dCas9-Control cells. This approach allowed us to assess the different levels of attachment based on which portions of the cell were in contact with the surface. Since stage 1 of attachment is unattached skimming, we omitted these cells from our analysis, and only included cells in stage 2 (lateral crest contact), stage 3 (lateral shield contact), and stage 4 (bare area contact). We found more cells at stages 2 and 3 and fewer in stage 4 for DAAP1-depleted cells than in the dCas9-Control (Figure 7B), indicating impaired seal formation around the ventral disc. Taken together, these data demonstrate that DAAP1 functions at the disc gates that regulate fluid flow and ventral disc seal formation.

## Discussion

Here, we have confirmed that *Gl*Actin has a role in ventral disc morphology and disc-based attachment (Figure 1). We also show that *Gl*Actin interactor DAAP1 is a stable component of the ventral disc (Figure 4). While perturbation of DAAP1 levels was not associated with visible effects on disc morphology, it resulted in decreased attachment (Figures 5–7).

The ventral disc is well known as a *Giardia-*specific microtubule based structure [17,32] and our data show that the actin cytoskeleton has an important role in ventral disc assembly and function. A functional relationship between actin and microtubules has been well-documented for other structures. In *Chlamydomonas*, actin is required for intraflagellar transport [33,34].

Actin monomers are a key component of the flagella through interaction with the inner dynein arm [35–38] and recent evidence indicates actin is a component of the gamma-tubulin ring complex [39]. Actomyosin interactions influence the orientation of the mitotic spindle [40,41] and F-actin associates directly with mitotic spindles in *Xenopus* and starfish embryos [42,43]. Actin is also involved in microtubule- or tubulin-based host interaction in other parasites. For instance, one of *Toxoplasma gondii’s* formins localize exclusively to the conoid [44], a latticelike organelle composed of tubulin which initiates host invasion [45]. There is evidence of coordination between the microtubules of *Toxoplasma*’s glideosome, which is used for actin-based motility, during host invasion [46]. Furthermore, actin polymerization is believed to contribute to conoid extrusion [47,48], and the conoid-localized Myosin H is essential for entry and exit from host cells [49,50]. *Gl*Actin polymerization could have an analogous role in controlling the movement of the ventral disc to facilitate attachment.

We also observed defects in the ventral disc in our *Gl*Actin*-*depleted cells (Figure 1), which we characterized as either mild (misshapen, broken, or with small gaps) or severe (unwound, severely misshapen, often associated with cytokinesis defects). There are several possible explanations for these phenotypes. During mitosis, the ventral disc is dismantled and two daughter discs are rebuilt during cytokinesis [27]. One possibility is that *Gl*Actin has a role in loading proteins on the assembling disc. *Gl*Actin also likely contributes to ventral disc positioning, since actin has a role in positioning basal bodies that nucleate the disc [51,52]. However, many of the cells with defective discs have normally-positioned flagella (Figure 1, Figure S2), indicating defective discs are not entirely due to basal body mispositioning. While the resolution of light microscopy limits our ability to see details in the microtubule array of the disc [53], over 25% of our unattached *Gl*Actin-depleted cells lacked noticeable defects (Figure 1). Therefore, a portion of the attachment defect cannot be attributed to disc defects alone.

Another possibility is that *Gl*Actin helps maintain the structure of the disc, consistent with its role in maintaining cell and organelle shape in other organisms [19,22]. The unusual stability of disc microtubules can be at least partially attributed to certain DAPs [8,15,17], which lock the microtubules into place, preventing their turnover. *Gl*Actin interacts with at least one of these DAPs, GL50803_5188 [18], and could be working in conjunction with it to stabilize the disc microtubules themselves. Another explanation is that *Gl*Actin works to stabilize one of the non-microtubule components of the ventral disc. The microtubules of the disc are overlaid by the giardin-composed microribbons [54], which are connected by crossbridges [13,55]. Since the spiral shape is often maintained, but unwound, in *Gl*Actin-depleted cells (Figure 1, Figure S2), it may be that *Gl*Actin contributes to establishing and/or maintaining disc overlap, as the overlap zone also functions as the anterior gate of the ventral disc.

A previous study from our lab identified eight *Gl*Actin interactors which localize to the ventral disc, including DAAP1 [18]; here we confirmed complex formation between *Gl*Actin and DAAP1 through co-immunoprecipitation (Figure 2). We did not detect co-localization between DAAP1 and *Gl*Actin in interphase cells, but present evidence that indicates co-localization during mitosis when new discs are assembled (Figure 2). Polymerized *Gl*Actin may be enriched in the ventral disc at these stages, but detection may be obscured due to technical limitations. Unfortunately, use of the anti-*Gl*Actin antibody does not differentiate between filaments and pools of monomeric *Gl*Actin which likely obscures some filaments. Currently, there are no methods to specifically detect F-*Gl*Actin, as Rhodamine-Phalloidin does not bind to *Gl*Actin filaments [22]. Actin in *Toxoplasma gondii* was believed to be primarily monomeric and only capable of forming short filaments [56,57]. However, expression of an anti-actin nanobody revealed a previously unseen complex network of F-actin [58]. Poor detection of *Gl*Actin using the anti-*Gl*Actin antibody could also be due to steric hindrance. Actin-binding proteins could sterically block access of anti-*Gl*Actin antibodies to disc-localized *Gl*Actin, similar to low anti-tubulin antibody staining compared to our ability to see microtubules of the disc with mNeonGreen-Tubulin [27]. Another possibility is that actin only transiently associates with the disc for specific conformational changes which would be challenging to capture in fixed cells. A live marker specific for *Gl*Actin is needed to determine the extent of actin association with conformational dynamics of the ventral disc including disc doming, bare area protrusion and disc flexion as seen in Movie S1.

Similar to other disc-associated proteins [44], DAAP1’s localization to the disc is stable, as we did not see any recovery of this protein for 12 minutes after photobleaching (Figure 4). While DAAP1 was previously identified as a potential disc interactor, it was not found in high enough abundance in the detergent-extracted disc proteome to qualify for further study as a disc-associated protein [14]. These results possibly indicating that this protein is a peripheral component of the disc, which would be expected in order for it to engage with a dynamic subset of *Gl*Actin.

Using CRISPR-interference, we depleted DAAP1 from cells, and found a mild defect in attachment, but no obvious impact on disc morphology (Figures 5–7). While a DAAP1 knockout would give us better insight into how DAAP1 loss affects the disc, the ability to readily knockout genes of interest remains under development. CRISPRi and Morpholino knockdown both have variable penetrance, and thus the level of depletion fluctuates among cells, and we observed normal discs even in cells with very little DAAP1 fluorescence (Figure 5). In contrast, depletion of median body protein with morpholinos results in broken discs [12]. Although an inability to attach can be attributed to disc defects [12,30], the depletion of gamma-giardin, another disc-localized protein that interacts with *Gl*Actin [18], resulted in malformed discs with normal attachment [59]. These findings indicate that some disc proteins may be involved in maintaining the structural integrity of the disc, while others are involved in controlling conformational changes of the disc required for modulating attachment. Our data is consistent with DAAP1 being in the latter category. [12]

The possibility that some disc proteins contribute to attachment but not structure may also explain the disparity between the severity of attachment defects in *Gl*Actin-depleted cells and DAAP1-depleted cells. Since many of the discs in *Gl*Actin-depleted cells were unwound or misshapen, this suggests roles in building the disc or collaboration with DAPs that maintain the disc structure. However, since over 25% of our unattached *Gl*Actin-depleted cells had apparently normal discs, the attachment defect cannot be attributed to disc defects alone. Our previous study identified eight other putative disc-localized F-*Gl*Actin interactors which could also contribute to attachment and/or disc morphology [18].

An additional explanation for the severity of attachment defects in *Gl*Actin-depleted cells is due to its involvements in supplemental mechanisms of attachment. *Gl*Actin has been demonstrated to function in building the ventrolateral flange, *Giardia’s* membrane lamella which also contributes to attachment [60,61]. Further, the marginal plate, a rigid structure associated with the anterior axonemes, is also hypothesized to be involved in attachment [11,32]. *Gl*Actin is stably enriched in this structure [22,62] and we have described six interactors which localize there as well [18]. Given *Gl*Actin’s involvement in all three structures hypothesized to augment attachment, depletion of *Gl*Actin likely has multivariant effects on attachment, contributing to a more severe phenotype.

The biology of parasites is often unlike anything seen in other organisms, shaped by the unusual environments in which they live. *Giardia’s* ventral disc exemplifies parasitic innovation and makes this *Giardia* specific structure a target for drug development. Since most DAPs lack homologs, they are in theory potential drug targets; however, the lack of active sites reduces their potential [63]. Further understanding of the mechanisms of attachment and its regulation will increase the viability of developing drugs that could specifically function through regulating attachment. Actin, however, is a druggable protein and conventional actin drugs do not bind to *Gl*Actin or inhibit its function, rendering it an appealing therapeutic target.

## Materials and Methods

### Parasite Strain and Growth Conditions

*G. lamblia* strain WB clone 6 (ATCC 50803; American Type Culture Collection) was cultured as described previously [22].

### Vector Construction

All constructs in this study were C-terminal fusions, made using Gibson reactions with linearized vectors and PCR products [64]. See Supplemental Table 3 for primer sequences and workflow.

### Live Cell Imaging

Cells were chilled with ice for 15 minutes to detach from the culture tube and then placed into an Attofluor cell chamber (Molecular Probes) and incubated in a GasPak EZ anaerobic pouch (BD) or a Tri-gas incubator (Panasonic) set to 2.5% O2, 5% CO_2_ for 90 minutes at 37°C. Cells were then washed four times with HEPES-buffered saline (HBS: 137mM NaCl, 5mM KCl, 0.91mM Na2HpO4-heptahydrate, 5.55mM Glucose, 20mM HEPES, pH7). Live cell imaging was performed on a DeltaVision Elite microscope (GE) equipped with DIC optics, using a 100 × 1.4 NA or 60 × 1.42 NA objective, and a sCMOS 5.4 PCle air-cooled camera (PCO-TECH).

### Coimmunoprecipitation

Cell cultures of 500 mL each of *Giardia* wildtype and DAAP1-3xHA were grown for 3 days, then iced for 2 hours to detach, and spun for 1500xg at 4°C for 20 minutes. Cells were then washed twice in 1X HBS with 2X HALT protease inhibitors and 10 µM chymostatin, 1 µM leupeptin, and 1 µM E64. Each pellet was resuspended to a final volume of 1.2 mL and 100 mM DSP in DMSO was added to a final concentration of 1 mM and incubated at room temperature for 30 minutes. The reaction was quenched for 15 minutes with an addition of Tris pH 7.4 final concentration 20 mM. Cells were then pelleted by spinning for 7 minutes at 700xg and resuspended in 350 µL lysis buffer (80 mM KCl, 10 mM imidazole, 1 mM MgCl2, 1 mM EGTA, 5% Glycerol, 20 mM HEPES, 0.2 mM, CaCl2, 10 mM ATP, 0.1% Triton X-100, 500 mM NaCl, pH 7.2, 2XHALT protease inhibitors and 10 µM chymostatin, 1µM leupeptin, and 1 µM E64). Cells were then lysed by sonication and cleared with a 10 minute spin at 10,000xg. A volume of 17.5 µL of equilibrated EZview Red Anti-HA Affinity gel (Sigma) was added to each tube of lysate, then incubated at 4 °C with end-over-end mixing for 1 hour. Beads were then spun at 8,200xg for 30 seconds and the supernatant was discarded, followed by a total of three washes with 750 µL wash buffer (80 mM KCl, 10 mM imidazole, 1 mM MgCl2, 1 mM EGTA, 5% Glycerol, 20 mM HEPES, 0.2 mM CaCl2, 10 mM ATP, 0.5% Tween, 500 mM NaCl, pH 7.2). Each wash consisted of end-over-end rotation for 5 minutes followed by a 30 second spin at 8,200 x g. Protein was then incubated with 50 µL of 8 M Urea at RT for 20 minutes to elute. The beads were pelleted and then sample buffer was added to the supernatant, the sample was boiled for 5 mins at 98 °C, and run on a 12% SDS-PAGE gel, followed by western blot protocol described previously [22].

### Immunofluorescence Microscopy

Fixation and imaging were performed as described in previous work [65], with the addition of a JaneliaFluor 646 HaloTag ligand (Promega, GA1120) or JaneliaFluor 549 HaloTag ligand (Promega, GA1110) at 1:100 in the second immunostaining step. Image analysis was performed in FIJI [66]. DAAP1 intensity was measured by generating a circular region of interest surrounding the ventral groove and measuring fluorescence intensity at the brightest point in each image stack.

### Morpholino Knockdown

Morpholino knockdown was performed as described previously [67]. Cells were lysed and run on a 10% SDS-PAGE gel, followed by western blot protocol as described previously [18,22]. For each of three biological replicates, samples were divided equally into three lanes and intensity of each lane measured using FIJI and averaged. Cell counts were performed with a MoxiZ coulter counter to ensure the same number of cells were included in each sample. For morpholino sequences, see Supplemental Table 3.

### Culture Tube Attachment Assay

*Gl*Actin: Cells were treated with either a standard control or anti-*Gl*Actin morpholino and allowed to recover for 24 hours in a total volume of 13 mL of TYDK media at 37 °C. At 24 hours, each culture tube was inverted six times; detached cells in media were decanted and counted with a MoxiZ coulter counter (Orflo), then replaced with 13 mL ice cold media. Culture tubes were placed on ice for 30 minutes to detach the remaining cells, after which these were also counted. Cells were then imaged live or fixed as described previously [27]. Three independent replicates of each experimental and control were analyzed. DAAP1 assays were performed similarly, except knockdown was accomplished with CRISPRi rather than morpholinos (see below).

### Assessment for Live/Dead Cells

Stock solutions of dye were created by dissolving 5 mg fluorescein diacetate (Sigma-Aldrich, F-7378) in 1 ml acetone and 2 mg propidium iodide (Sigma-Aldrich, P4170) in 1 ml HBS. A 2X stain solution was prepared by diluting the stock solutions of fluorescein diacetate and propidium iodide by 1:100 in 1X HBS. Cells were pelleted by centrifugation at 500 x g and resuspended in 1 mL 1X HBS. 1mL of 2x stain solution was then added to the cell suspension, washed twice in 1X HBS by centrifugation and resuspended in 1X HBS for imaging.

### Fluorescence Recovery after Photobleaching

FRAP was performed on DAAP1-mNeonGreen in the ventral disc to measure protein turnover. Cells were chilled with ice for 20 minutes to detach from the culture tube and then placed into an Attofluor cell chamber (Molecular Probes) and incubated in a O_2_/CO_2_ incubator (Panasonic) for 1-2 hours at 37° C. Media was replaced with 37 °C 1X HBS before imaging. All experiments were conducted using a FRAP-enabled DeltaVision Spectris with 405 nm solid-state laser, 100% laser power, 50 mW laser, and 25 ms stationary pulse. Three images were taken until the bleach event for 0.5 second intervals, followed by image acquisition every 10 sec for 10 minutes. Normalized mNeonGreen fluorescence recovery was calculated by subtracting the background noise from the ROI intensity measurement; to normalize for photobleaching due to imaging, this measurement was then divided by a fluorescent control ROI intensity measurement with the background noise subtracted as well.

### CRISPR Interference Knockdown

Constructs were designed as described previously [68]. For primer design and workflow, see Supplemental Table 3. Cells containing an integrated C-terminally mNeonGreen-tagged DAAP1 were transfected with a dead Cas9 vector which contained either a non-specific small guide RNA (dCas9-CTRL) or a small guide RNA targeting base pairs 392 through 411 of the coding region of DAAP1 (dCas9+sgRNA392-411). Knockdown efficiency was measured using a Tecan Spark plate reader. Cells were pelleted and washed with 1X PBS and then resuspended in 1X PBS. An equal number of dCas9-CTRL or dCas9+sgRNA392-411cells were loaded into Tecan black flat 96-well plates and the volume was adjusted to a total of 150 µL. Three measurements per sample were averaged and three independent transformations were averaged for total knockdown efficiency, normalized to the control and blanked with 1X PBS.

### Shear Force Attachment Assay

Microfluidic devices were fabricated using a silicon master, which was created using deep reactive ion etching. The silicon master contained a raised channel that had a rectangular cross-sectional area, with a length of 1 cm, a width of 1 mm, and a height of 115 µm. A polydimethylsiloxane (PDMS) (Sylgard 184, Dow Corning) mold was made using a 10:1 base to curing agent ratio, which was mixed for 5 minutes, degassed for 20 minutes, poured over the master, and placed in an oven for 10 minutes at 110 °C to cure. The mold was peeled away from the silicon master and holes were punched for the inlet and outlet of the channel. Afterwards, inlet and outlet ports were created using a custom aluminum mold with pegs to insert the silicone tubing (0.040 inch ID/ 0.085 inch OD, HelixMark). Degassed 10:1 PDMS was poured into the preheated aluminum mold and cured at 110 ◦C for 1 hour. The inlet and outlet ports were aligned with holes on the top face of the microfluidic device and stuck together using uncured PDMS and 5 minutes of oven heat. Finally, the bottom face of the microfluidic device and a clean glass slide were plasma treated for 10 seconds and aligned together to create a water-tight seal between the two layers.

Control dCas9 or DAAP1 targeting dCas9+sgRNA392-411 trophozoites were chilled for 20 minutes on ice and then loaded into microfluidic devices placed in a GasPak EZ anaerobic pouch (BD Scientific) at 37 °C for 1 h to allow re-attachment. Growth media was drawn into a disposable syringe (BD Scientific) and then loaded onto a syringe pump (Harvard Apparatus). For experiments, the flow rate was set to 20 µl/min for 30 s to clear swimming trophozoites and then ramped to 100 µl/min over 5 s and challenged for 60 s. The challenge assays were recorded at 2 FPS using a Nikon Eclipse Ti microscope with a 40X objective, using phase illumination, Clara DR-1357 CCD camera (Andor), and a live-cell incubation chamber (In Vivo Scientific, Inc) to maintain a temperature of 37 °C. Trophozoite mean speed was quantified starting at 35s using the TrackMate ImageJ plugin via manual tracking over the first 10 seconds of 100µL/min flow [31].

### TIRF Microscopy

Cells were prepared for live imaging as described above, with the addition of CellMask™ Orange Plasma membrane Stain (ThermoFisher) according to manufacturer’s instructions after media replacement with 1X HBS. Cells were imaged in a chamber with 2% CO2 and 5% O2 with a DeltaVision OMX (GE) using 60X TIRF objective. Images were taken from randomly selected fields of view. Each dish was imaged for a maximum of 30 minutes to minimize stress on the cells. Three dishes from two independent experiments were used for each condition.

### Live Imaging with Microspheres

Cells expressing DAAP1-mNeonGreen were transfected with a plasmid containing Halo-tagged delta-giardin and either dCas9 control or dCas9+sgRNA392-411 to deplete DAAP1. Cells were grown in parallel and upon confluency were chilled with ice for 15 minutes to detach from the culture tube and then placed into an Attofluor cell chamber (Molecular Probes), then incubated in a GasPak EZ anaerobic pouch (BD) or a Tri-gas incubator (Panasonic) set to 2.5% O2, 5% CO_2_ for 90 minutes at 37 °C. 646 HaloTag ligand (Promega, GA1120) was added to cells 45 minutes after incubation at 1:1000 concentration, incubated for 15 minutes, and then replaced with TYDK for 30 minutes prior to imaging. 2.5 µL of fluorescent microspheres (0.2 µm, Polysciences Fluoresbrite® YO Carboxylate Microspheres catalog number 19391) were added to 0.5 mL 1xHBS and sonicated for 10 minutes using a bath sonicator to disrupt aggregates. This diluted solution was then added to cells at a 1:10 ratio to each chamber (30 µL diluted bead solution in 300 µL cells). Images of single focal planes in which the disc was visible were taken every 1 second for 15 seconds for each condition. Cells with beads which were collected under their ventral disc and cells with no beads were counted. Cells which detached or moved were excluded from analysis. Live cell imaging was performed on a DeltaVision Elite microscope (GE) equipped with DIC optics, using a 60 × 1.42 NA objective, and a sCMOS 5.4 PCle air-cooled camera (PCO-TECH).

## Acknowledgements

We thank Kelly Hennessey, Kelli Hvorecny, Han-wei Shih, Elizabeth Thomas, Barbara Wakimoto, Susan Parkhurst and Germain Alas for manuscript editing. We thank Pang Chan for assistance with TIRF microscopy at the Biology Imaging Facility.

This material is based upon work supported by the National Science Foundation Graduate Research Fellowship under Grant No. DGE-1762114 to MSO. Funding for the OMX was provided by grant proposal S10 OD021490. Research reported in this publication was supported by the National Heart, Lung, And Blood Institute of the National Institutes of Health under Award Number F31HL156697. The content is solely the responsibility of the authors and does not necessarily represent the official views of the National Institutes of Health.”

---

**Supplemental figure 1: *Gl*Actin Depletion by antisense translation-blocking Morpholinos A)** Representative western blot of cells treated with either a standard control morpholino (left) or morpholino targeting *GlActin* (right) for 24-hours and probed for tubulin (cyan) and *Gl*Actin (magenta), resulting in a ~61% reduction of *Gl*Actin levels. **B)** Average of three morpholino knockdowns results in an average of 50.3% ± 5.6 reduction of protein expression, as determined by western blot. Values shown are mean ± SEM.

**Supplemental Figure 2: Range of Disc Defect Phenotypes in *Gl*Actin-Depleted Cells A)** Representative *Giardia* cell from the control (attached) condition. **(B-E**) are *Gl*Actin depleted cells with disc defects. Mild defects were categorized as discs which were broken **(B)** or misshapen **(C)**. Severe defects included unwound discs (**D**) or disc defects associated with failed cytokinesis **(E)**. All cells pictured in B-E were unattached, with the exception of the bottom row of B. Scale bar denotes 5 µm.

**Supplemental Table 1:** Raw number of *Gl*Actin-depleted attachment assay cell counts and statistics

**Supplemental Table 2: DAAP1-depleted attachment assay cell counts and statistics**

**Supplemental Table 3: Primer and morpholino sequences and workflow**

**Supplemental Table 4: Number of microspheres per cell and statistics**

