## Supplementary figures and images for "Disc and Actin-Associated Protein 1 Influence Attachment in the Intestinal Parasite *Giardia lamblia*"

### Figure S1

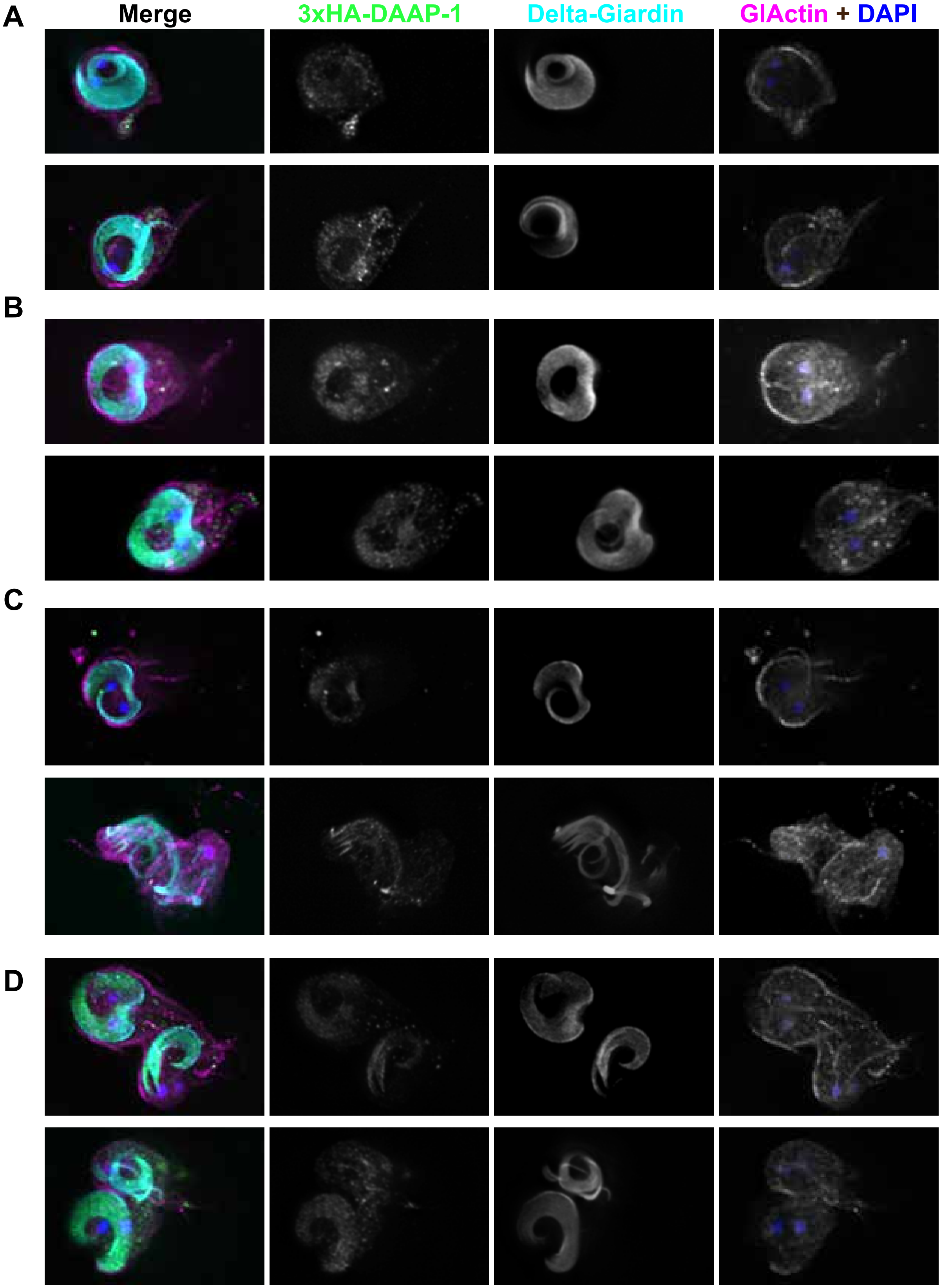

### Figure S2

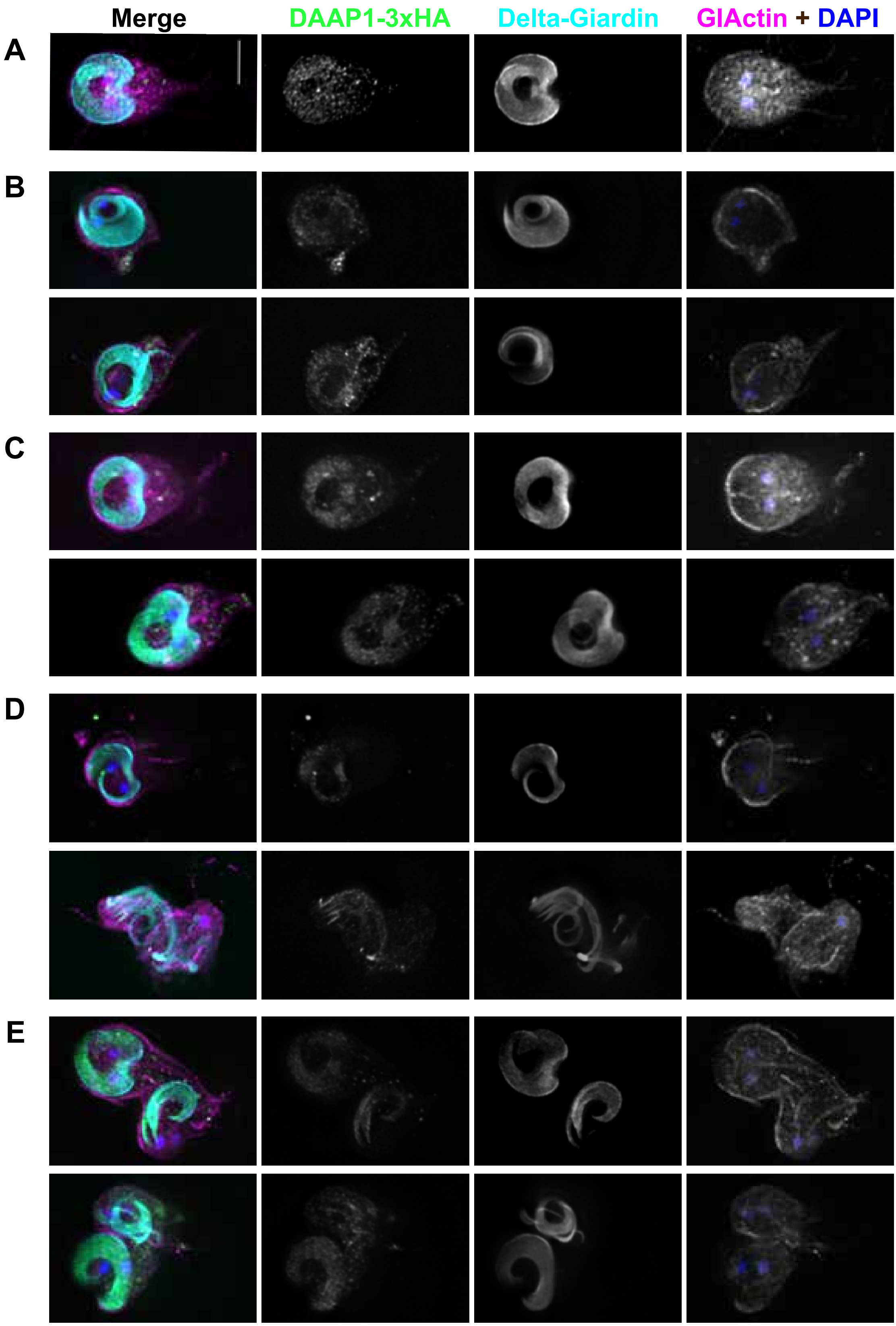
